# Multiobjective Strain Design: A Framework for Modular Cell Engineering

**DOI:** 10.1101/349399

**Authors:** Sergio Garcia, Cong T. Trinh

## Abstract

Diversity of cellular metabolism can be harnessed to produce a large space of molecules. However, development of optimal strains with high product titers, rates, and yields required for industrial production is laborious and expensive. To accelerate the strain engineering process, we have recently introduced a modular cell design concept that enables rapid generation of optimal production strains by systematically assembling a modular cell with an exchangeable production module(s) to produce target molecules efficiently. In this study, we formulated the modular cell design concept as a general multiobjective optimization problem with flexible design objectives derived from mass action. We developed algorithms and an associated software package, named ModCell2 to implement the design. We demonstrated that ModCell2 can systematically identify genetic modifications to design modular cells that can couple with a variety of production modules and exhibit a minimal tradeoff among modularity, performance, and robustness. Analysis of the modular cell designs revealed both intuitive and complex metabolic architectures enabling modular production of these molecules. We envision ModCell2 provides a powerful tool to guide modular cell engineering and sheds light on modular design principles of biological systems.

## INTRODUCTION

Engineering microbial cells to produce bulk and specialty chemicals from renewable and sustainable feedstocks is becoming a feasible alternative to traditional chemical methods that rely on petroleum feedstocks (1). However, only a handful of chemicals, out of the many possible molecules offered by nature, are industrially produced by microbial conversion, mainly because the current strain engineering process is laborious and expensive for profitable biochemical production (2). Thus, innovative technologies to enable rapid and economical strain engineering are needed to harness a large space of industrially-relevant molecules (3).

The modular organization of biological systems has been a source of inspiration for synthetic biology and metabolic engineering (4, 5). Modular pathway engineering breaks down target pathways into tractable pathway modules that can be finely tuned for optimal production of desirable chemicals (6, 7). Harnessing combinatorial pathways (e.g., fatty acid biosynthesis, reverse beta oxidation, polyketide or isoprenoid biosynthesis) is one excellent example of modular pathway engineering. These pathways contain metabolic similarity (or combinatorial characteristics) such as a group of common specific enzymes capable of catalyzing linear reaction steps (8) and/or elongation cycles (9-11) and hence are capable of producing a large library of unique molecules (12). Since these molecules are derived from a common precursor metabolite(s), the optimal production strains often share common genotypes and phenotypes, and hence, the costly strain optimization process is only performed once for these molecules. Remarkably, this advantageous strain optimization strategy can be applied even for production of molecules derived from different precursors, using the concept of modular cell (ModCell) design (2, 13, 14).

With the arrival of steady-state, constraint-based stoichiometric models of cellular metabolism, various computational algorithms have been developed to guide strain engineering (15-17). These methods have featured the design of strains capable of growth-coupled product synthesis (*GCP*), enabling adaptive laboratory evolution of these designed strains to enhance product titers, rates, and yields (14, 18-20). Two approaches on growth-coupled production have been formulated — one based on the coexistence of maximum growth and product synthesis rates during the growth phase (21) and the other based on the obligate requirement of optimal product synthesis in any growth phase (22). The distinction between these two types of growth coupling are also referred to weak coupling (*wGCP*) and strong coupling (*sGCP*) (16, 23).

Development of most strain design algorithms has been focused on overproduction of only one target molecule. The first algorithm proposed for modular cell design compatible for overproduction of multiple target molecules is MODCELL (13), which guided several experimental studies (14, 24-27). It works by generating *sGCP* strain designs for each target product based on elementary mode analysis (28), and then comparing the design strategies of different products to identify common genetic modifications among them. A similar approach was adapted in a subsequent work (29). For MODCELL to find optimal solutions for multiple target products, it requires: 1) enumerating all possible designs above a predefined minimum product yield and with minimal reaction deletion sets for each production network, which might lead to a large number of solutions for each network and hence make the problem computationally intractable, and 2) the resulting designs for all products must be compared to identify common interventions, which is a computationally-hard, set-covering problem. Thus, the current enumerative approach of MODCELL might become intractable very quickly, especially for large-scale metabolic networks and potentially generate non-optimal designs, i.e., requiring more knockouts than necessary or including fewer products than possible.

In this study, we generalized the concept of modular cell design and addressed the computational limitation of implementing it. We developed a novel computational platform (ModCell2), based on multiobjective optimization and analysis of mass action of cellular metabolism, to guide the design of modular cells for large-scale metabolic networks. We demonstrated that ModCell2 can systematically identify genetic modifications to design modular cells that can couple with a variety of production modules and exhibit a minimal tradeoff among modularity, performance, and robustness. By analyzing these designs, we further revealed both intuitive and complex metabolic architectures enabling modularity in modular cell and production modules required for efficient biosynthesis of target molecules.

## METHODS

### Design principles of modular cell engineering

In the conventional strain engineering approach, a parent strain is genetically modified to yield an optimal production strain to make only a target product. To produce each new molecule, the design-build-test cycles of strain engineering must be repeated, which is laborious and expensive (Figure 1). To minimize the cycles, modular cell engineering is formulated by genetically transforming a parent strain into a modular (chassis) cell that must be assembled with exchangeable modules to create optimal production strains (13). A modular cell is designed to contain core metabolic pathways shared across designed optimal production strains. Exchangeable modules are production pathways designed to synthesize desirable chemicals. A combination of a modular cell and a production module(s) is required to balance redox, energy, and precursor metabolites for sustaining cellular metabolism during growth and/or stationary phases and exhibiting only desirable phenotypes. Practically, modular cell engineering can be applied to monocultures and polycultures, where a production module(s) can be embedded in a modular cell and activated by intracellular and/or extracellular cues such as light and/or signaling molecules.

**Figure 1:**
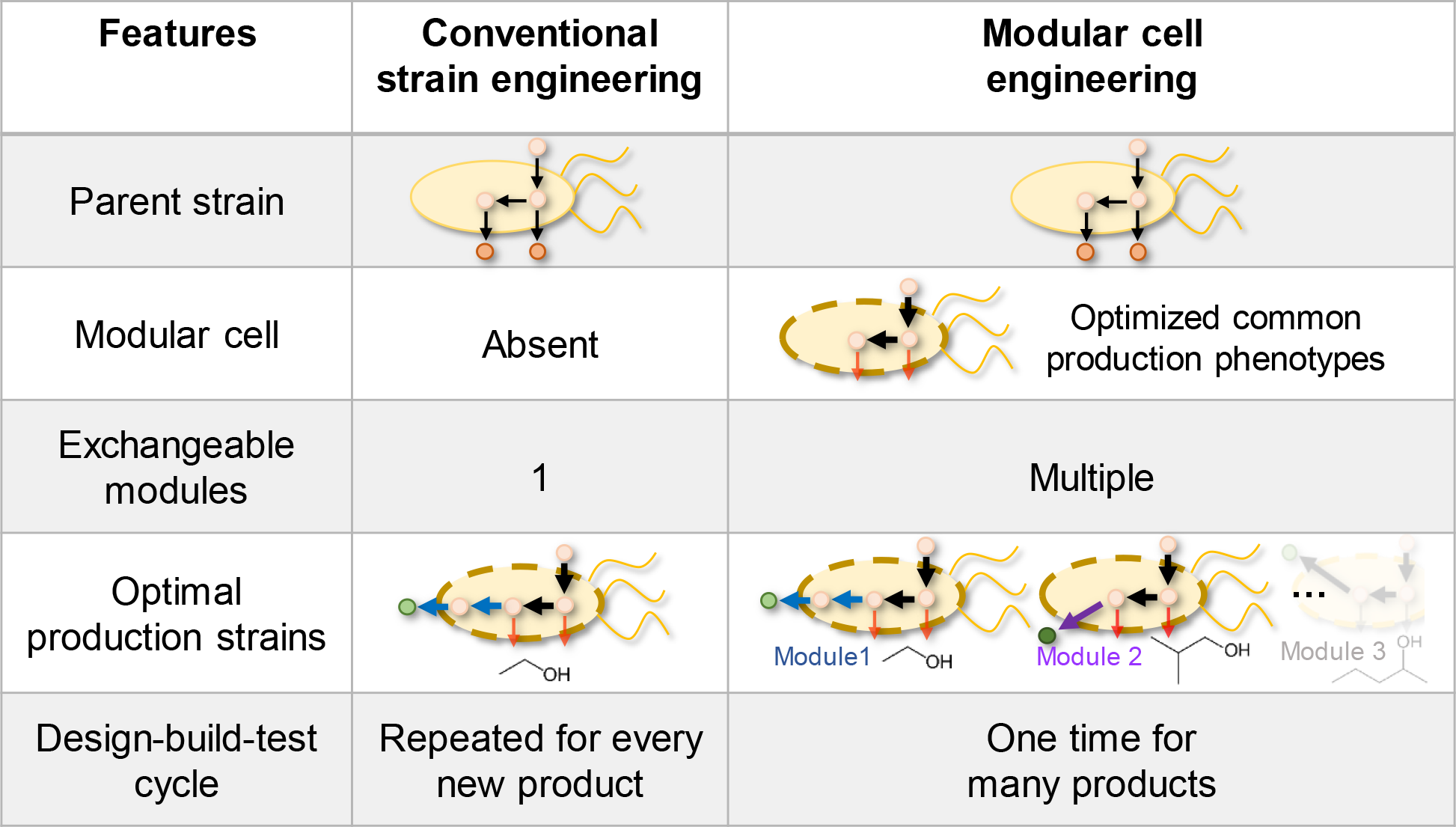
Comparison between the conventional single-product strain design and modular cell engineering. In the conventional approach, each target product requires to go through the iterative optimization cycle. The modular cell engineering approach exploits common phenotypes associated with high product titers, rates, and yields; and hence, the strain optimization cycle only needs to be performed once for multiple products, which helps reduce the cost and time of strain development.

### Multiobjective strain design framework for modular cell engineering

For modular cell engineering, we seek to design a chassis cell compatible with as many production modules as possible to achieve only desirable production phenotypes while requiring minimal genetic modifications. Since all production modules must leverage cellular resources of the modular cell (e.g. precursor metabolites, cofactors, and energy), they form competing objectives. Therefore, the framework of modular cell engineering can be formulated as a multiobjective optimization problem, named ModCell2, as described below.

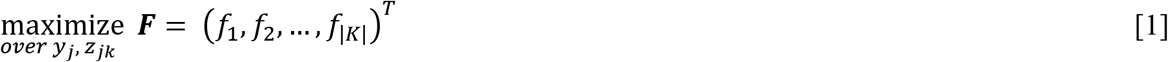

*subject to*

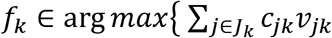

*subject to*

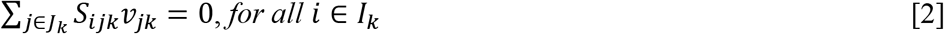

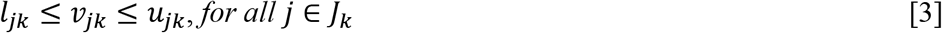

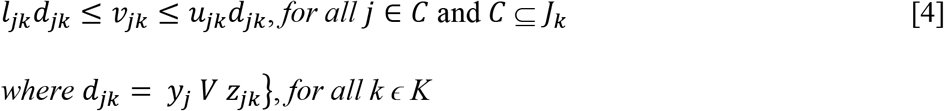

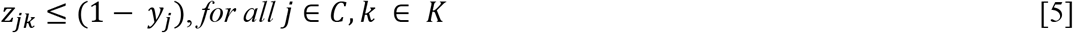

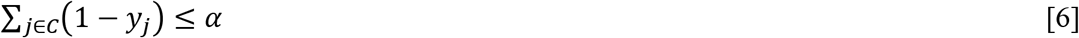

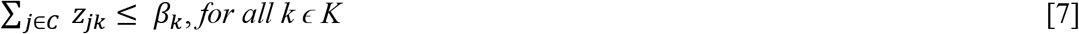

where *i*, *j*, and *k* are indices of metabolite *i*, reaction *j*, and production network *k*, respectively; *f*_*k*_ is a design objective for network *k*; *c*_*jk*_ represents the cellular objective for reaction *j* in network *k* associated with a design objective defined in eqns. 8-10; *v*_*jk*_ (mmol/g DCW/h) is metabolic flux of reaction *j* bounded by *l*_*jk*_ and *u*_*jk*_ in network *k*, respectively; *y*_*j*_ and *z*_*jk*_ are binary design variables for deletion reaction *j* and module reaction *j* in network *k*, respectively; *α* and *β*_*k*_ are design parameters for deletion and module reactions, respectively; *S*_*ijk*_ is a stoichiometric coefficient of metabolite in reaction *j* of network *k*; and *C* (eqn. 4) is the candidate reaction set (Supplementary File S1). The goal of the optimization problem is to simultaneously maximize all design objectives *f*_*k*_.

#### Steady-state mass balance constraint of cellular metabolism

Quasi steady-state flux balance of cellular metabolism (eqn. 2) is used as metabolic constraints for eqn. 1 (30, 31). A model corresponding to each modular production strain (i.e. production network k) will be derived from a parent strain (i.e. parent network) by adding necessary reactions (e.g., a production module) to produce a target molecule. A feasible flux distribution for each production network is described by mass balance (eqn. 2) and reaction flux bounds (eqns. 3 and 4). For a given production network, the phenotypic space can be illustrated by the gray area that is projected onto the two-dimensional space spanned by product synthesis and growth rates (Figure 2).

**Figure 2:**
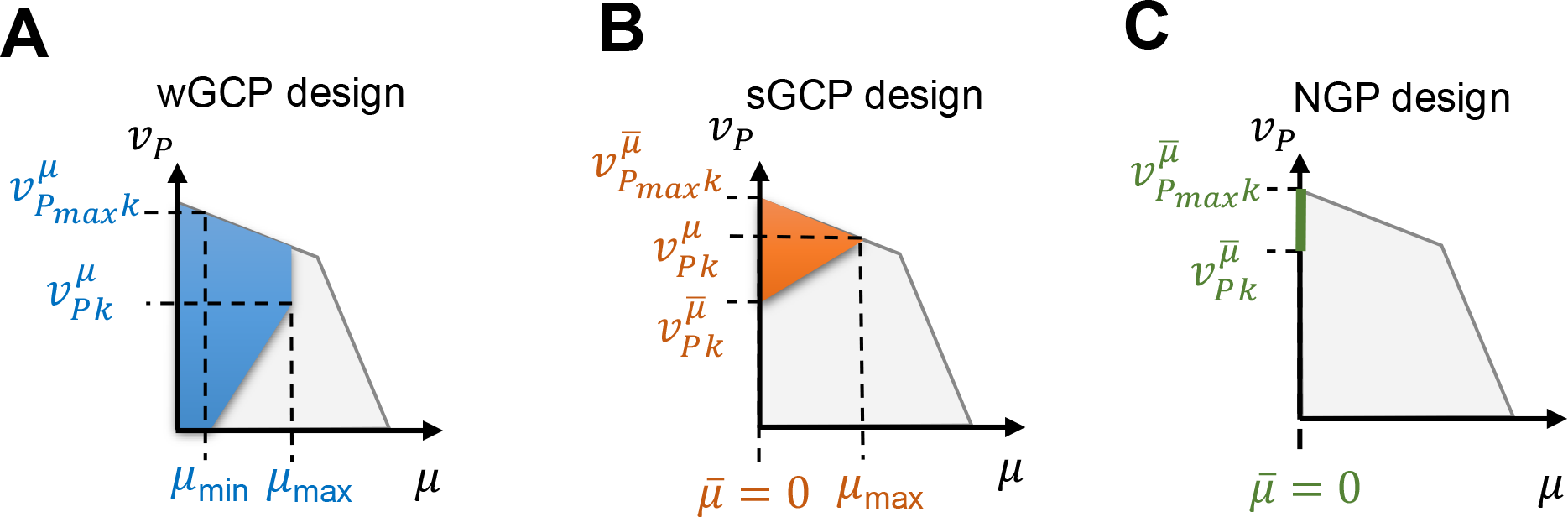
Graphical representation of phenotypic spaces for different strain design objectives including **(A)** weak growth coupling (wGCP), **(B)** strong growth coupling (*sGCP*), and **(C)** nogrowth production (*NPG*). 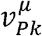 is the minimum product formation rate at the maximum growth rate for production network *k*, and 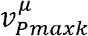 is the maximum product secretion rate attainable. 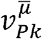 and r!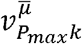 are the minimum and maximum product formation rates for production network *k* during the stationary phase, respectively.

#### Design variables

In our formulation for modular cell engineering, we introduced two design variables - binary reaction deletions (*y*_*j*_) inherent to the modular cell and module-specific reaction insertions (*z*_*jk*_) (eqn. 4). These variables can be experimentally manipulated to constrain the desirable phenotypes of production strains as shown in Figure 2. Specifically, *y*_*j*_ = 0 if reaction *j* is deleted from the modular cell; otherwise, *y*_*j*_ = 1. Deleting metabolic reactions removes undesired functional states of the network and leaves those with high design objectives. Likewise, *z*_*jk*_ = 1 if reaction *j* is present in the production network *k*; otherwise, *z*_*jk*_ = 0. These module reactions are endogenous reactions removed from the parent network (eqn. 5), but are added back to a specific production module to enhance the compatibility of a modular cell. The maximum number of reaction deletions (*α*) and module-specific reaction insertions (*β*_*k*_) are user-defined parameters.

#### Design objectives

To generalize ModCell2 design, we allow three different types of design objectives (*f*_*k*_, eqn. 1) that determine production phenotypes for each production network. Depending on the application, a phenotype can be designed to be weak coupling (*wGCP*), strong coupling (*sGCP*), and/or non-growth production (*NGP*) (Figure 2). The constrained phenotypic spaces based on these design objectives are shown in color; any point within these spaces is a feasible physiological state of the cell that can be represented by a metabolic flux distribution.

The *wGCP* design seeks to achieve a high product rate at maximum growth rate (Figure 2A). The *wGCP* design objective, 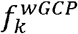 (∈[0,1]), is calculated as follows:

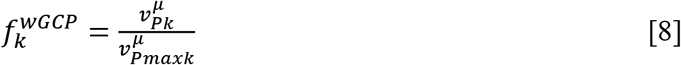

where 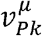 is the minimum synthesis rate of the target product P at the maximum growth rate for production network *k* and 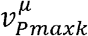. is the maximum synthesis rate of P (Supplementary File S1). This *wGCP* design formulation is equivalent to RobustKnock (32) or OptKnock with a tilted objective function (21, 33, 34). In eqn. 8, 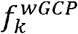 is scaled from 0 to 1 for proper comparison among products. The *wGCP* design is appropriate for applications where growth rate is not limited by the nutrients, and the product is formed during the growth phase.

The *sGCP* design seeks to achieve a high product rate not only at optimal growth rate but also during non-growth phase (Figure 2B). The *sGCP* design objective, 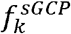, (∈[0,1]), is calculated as follows:

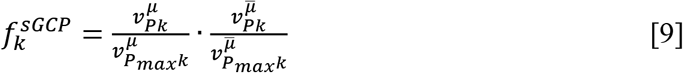

where 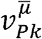 and 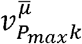 are the minimum and maximum product formation rates for production network *k* in the stationary phase, respectively (Supplementary File S1). The *sGCP* design objective is comparable to the one implemented in MODCELL (13). Different from *wGCP*, *sGCP* requires high product synthesis rate for any growth phase. However, the additional constraint of optimal product synthesis during the stationary phase requires more genetic manipulations or specific experimental conditions (e.g., anaerobic growth condition, supply of intermediate metabolites). Both *wGCP* and *sGCP* designs enable fast growth selection to attain the optimum product rates by adaptive laboratory evolution (35, 36).

The *NGP* design aims to maximize the minimum product rate during the non-growth phase by eliminating carbon fluxes directed to biomass synthesis (Figure 2C). The *NGP* design objective, 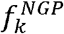 (∈[0, 1]), is calculated as follows:

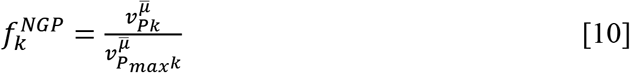

While the *NGP* design is not suitable for growth selection, it can be derived from a *wGCP* (or *sGCP*) design by imposing additional genetic modifications. Practically, *NGP* design strains can be activated during cell culturing using a regulatory genetic circuit to toggle switch between production phases.

#### Design solutions

Optimal solutions for eqn. 1 are a Pareto set (***PS***) that correspond to design variables, including reaction deletions (*y*_*j*_) and module reaction insertions (*z*_*jk*_). Each solution constitutes a design of a modular cell:

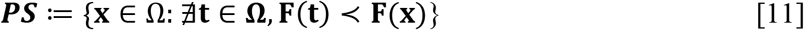

Here, **F(t)** ≺ **F(x)** means **F(t)** *dominates* **F(x)** if and only if f_i_(**t**) ≥ f_i_(**x**) for all *i*, and **F(t)** differs from **F(x)** in at least one entry. The feasible space of design variables, Ω, is defined by the problem constraints (eqn. 2-7, also see Supplementary File S1). Phenotypes of modular cells will be the image of the Pareto set in the objective space, known as the Pareto front (***PF***):

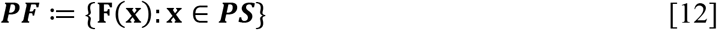

For the multiobjective strain design framework, the input parameters include *α* (eqn. 6), *β*_*k*_ (eqn. 7), and the production networks as input metabolic models. Each model contains a production module to produce one target chemical. The output is a Pareto set (genetic modifications) and its respective Pareto front (desirable production phenotypes). For a special case with no trade-off among the design objectives, an optimal solution, named a utopia point, exists where each objective achieves its maximum value. The multiobjective strain design formulation presented is general and can be applied to design modular cells for any organism.

### Algorithm and implementation

#### ModCell2 algorithm

To solve the multiobjective optimization problem for modular cell engineering, we used multiobjective evolutionary algorithms (MOEAs) (37). MOEAs were selected because they can efficiently handle linear and non-linear problems and do not require preferential specification of design objectives (38). MOEAs start by randomly generating a population of individuals (a vector of design variables), each of which is mapped to a design objective vector (i.e., a fitness vector). In ModCell2 (Supplementary File S1), the objective values of an individual are calculated by solving the linear programming problems for each production network. Next, individuals are shuffled to generate an offspring, from which the most fit individuals are kept. This process was repeated until the termination criteria was reached, for instance, either the solutions cannot be further improved or the simulation time limit is reached.

#### ModCell2 implementation

To streamline the modular cell design, we developed the ModCell2 software package based on three core classes (Figure S1 in Supplementary File S2). The Prodnet class parses and pre-processes production network models, and computes production phenotypes. The MCdesign class serves as an interface between the MOEA optimization method and metabolic models. Finally, the ResAnalysis class loads the Pareto set computed by MCdesign and identifies the most promising modular cell designs.

The code was written in MATLAB 2017b (The Mathworks Inc.) using the function gamultiobj() from the MATLAB Optimization Toolbox that implements the NSGA-II algorithm (39) to solve the multiobjective optimization problem. The solution and analysis methods were parallelized using the MATLAB Parallel Computing Toolbox. The linear programs to calculate metabolic fluxes were solved using the GNU Linear Programming Kit (GLPK). The COBRA toolbox (40, 41) and F2C2 0.95b (42) were also used for COBRA model preprocessing and manipulation.

#### Metabolic models

In our study, we used three parent models including i) a small metabolic network to illustrate the modular cell design concept (13), ii) a core metabolic network of *Escherichia coli* to compare the performance of ModCell2 with respect to the conventional single-product strain design strategy and the first-generation modular cell design platform MODCELL (13), and iii) a genome-scale metabolic network of *E. coli* (i.e., iML1515 (33)) for biosynthesis of a library of endogenous and heterologous metabolites, including 4 organic acids, 6 alcohols, and 10 esters (Figures S2 in Supplementary File S2) (8, 22, 25, 43-49).

#### Simulation protocols

Anaerobic conditions were imposed by setting oxygen exchange fluxes to be 0, and the glucose uptake rate was constrained to be at most 10 mmol/gCDW/h, as experimentally observed for *E. coli.* When using the genome-scale model iML1515 to simulate *wGCP* designs, the commonly observed fermentative products (acetate, CO2, ethanol, formate, lactate, succinate) were allowed for secretion as described elsewhere (50). For simulation of *sGCP* and *NGP* designs, the glucose uptake rate was fixed (i.e., −10 mmol/gCDW/h); otherwise, the flux is not active during the no-growth phases, resulting in the product synthesis rate of 0 regardless of genetic manipulations. To compare ModCell2 with Optknock, we applied the OptKnock algorithm with a titlted objective function (51) to generate *wGCP* designs for each production network, using the open-source algebraic modeling language Pyomo (52). The MILP problems were solved using CPLEX 12.8.0 with a time limit of 10,000 seconds set for each product. ModCell2 is provided as an open-source software package and is freely available for academic research. The software package and documentation can be download via either web.utk.edu/~ctrinh or Github https://github.com/TrinhLab.

### Analysis methods for design solutions

#### Compatibility

The compatibility, *C* (ϵ *Z*+), of a design is defined as the number of products that are coupled with a modular cell and has objective values above a specified cutoff value *θ* As a default, we set *θ* = 0.6 for the *wGCP* and *NGP* design objectives and *θ* = 0.36 (0.6^2^) for the *sGCP* design objective. For example, a *wGCP* design for 3 products that has the design objective values of 0.4, 0.9, and 0.6 has a compatibility of 2, given a cutoff value of *θ*≥ 0.6.

#### Compatibility difference and loss

Robustness is the ability of a system to maintain its function against perturbations, and hence is very important of designing biological and engineered systems (53). To evaluate the robustness of modular cell designs, we defined two metrics, the compatibility difference (*CD*) and compatibility loss (CL ϵ [0,1]) as follows:

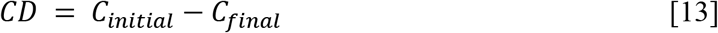

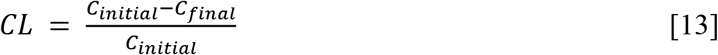

where *C*_*initial*_ and *C*_*final*_ are the compatibilities of a modular cell design before and after a single reaction deletion, respectively. The value *CD* > 0 (or *CL* > 0) means the modular gains fitness while *CD* < 0 (or *CL* < 0) means that it loses its fitness. In the analysis, we did not consider essential and blocked reactions for our single-deletion analysis; for instance, there are only 1139 potential reaction deletions in the iML1515 model.

#### Metabolic switch design

A metabolic switch design is a modular cell that can possess multiple production phenotypes (i.e., *wGCP*, *sGCP*, and *NPG*), activated by an environmental stimulus (e.g. metabolites, lights). The metabolic switch design is enforced to have a set of reaction (gene) deletions in one production phenotype to be a subset of the other, for instance, {**y**_wgcp_} ⊆ {**y**_NPG_}, {**y**_wWep_} ⊆ {**y**_sGCp_}. The metabolic switch design is beneficial for multiphase fermentation configurations that enable flexible genetic modification and implementation. Specifically, the metabolic switch design can exhibit the *wGCP* phenotype during the growth phase and the *NPG* (or *sGCP*) phenotype during the stationary phase. The metabolic switches can be implemented using the genetic switchboard (54).

## RESULTS AND DISCUSSION

### Illustrating ModCell2 for modular cell design of a simplified network

An example parent network, adapted from (13), was used to illustrate ModCell2 (Figure 3A). Inputs for the multiobjective optimization problem include i) three production networks (Figure 3B), comprising of one endogenous production module (module 1) and two heterologous production modules (modules 2 and 3) and ii) design parameters (Figure 3C), containing design objective type, maximum number of deletion reactions (*α*), and maximum number of module reactions (*β*_*k*_). The output of ModCell2 generated the Pareto set and the corresponding Pareto front for modular cell designs (Figure 3D). The 2-D plots of product yields versus growth rates presented the feasible phenotypic spaces of the wildtype (gray area) and the designed strain (blue area).

**Figure 3:**
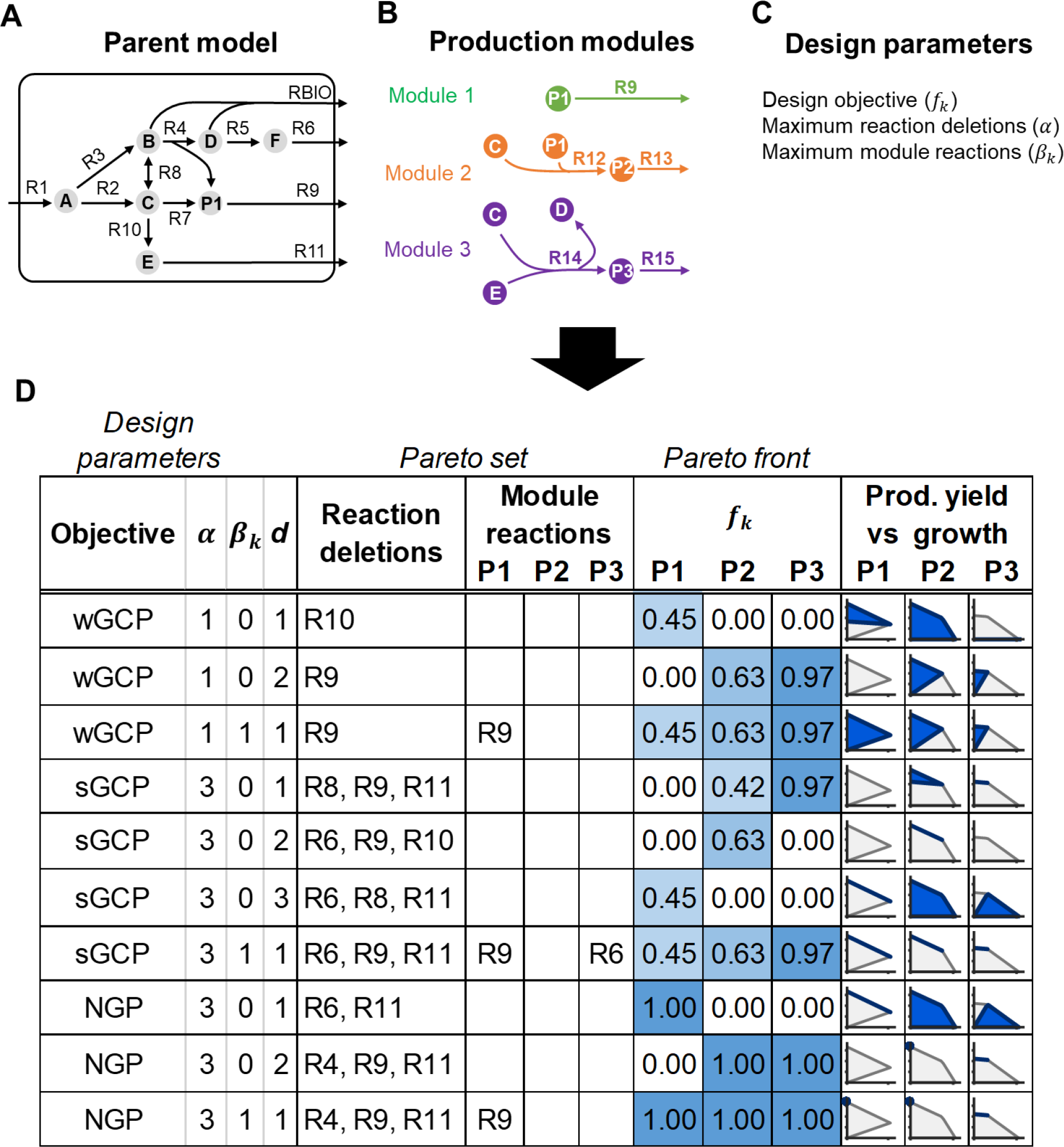
Illustration of ModCell2 workflow and analysis including **(A)** parent model, **(B)** production modules, **(C)** design parameters, and **(D)** simulation output for Pareto set and Pareto front based on design input.

Using various *α* and *β*_*k*_ values, ModCell2 simulation generated three *wGCP*-*α*-*β*_*k*_-d, four *sGCP*-*α*-*β*_*k*_-d designs, and three *NGP*-*α*-*β*_*k*_-d designs, where *d* is the design solution index (Figure 3D). For instance, by setting *α* = 3 and *β*_*k*_ = 0, we found three *sGCP* designs including *sGCP-3-0-1*, *sGCP-3-0-2*, and *sGCP-3-0-3.* The first design *sGCP-3-0-1* has a compatibility of 2 (given θ > 0) with the design objective values of 0.42 and 0.97 for the products P2 and P3, respectively. In contrast, the *sGCP-3-0-2* and *sGCP-3-0-3* designs have compatibilities of only 1 with the design objectives of 0.63 for P2 and 0.45 for P1, respectively.

Based on all designs, we can clearly see the trade-offs for optimization of different products for *β*_*k*_ = 0. However, setting *β*_*k*_ ≥ 1 helps increase the compatibility of a modular cell with different production modules. In addition, we found that the Pareto front collapses into a utopia point as seen in the *wGCP-1-1-1*, *sGCP-3-1-1*, and *NGP-3-1-1* designs. For instance, the modular cell, *sGCP-3-1-1*, is compatible with all three products. The three corresponding optimal production strains can couple growth and product formation during the growth phase. During the stationary phase, these strains produce the products at maximum theoretical yields. In theory, a universal modular cell always exists, provided that enough reaction deletions and module reactions are used. It might be more tractable to construct such a modular cell from a synthetic minimal cell using the bottom-up approach. However, construction of a universal modular cell from a host organism (e.g., *E. coli, S. cerevisiae*) using the top-down approach will require a significantly large number of genetic modifications, that might be challenging.

### Comparing ModCell2 designs with first-generation MODCELL and single product designs

#### ModCell2 can generate more and better designs than the first-generation modular cell design platform

To evaluate the algorithms and performance of ModCell2, we directly compared it with MODCELL (13) in two case studies, using the same core model of *E. coli* for production of five alcohols (ethanol, propanol, isopropanol, butanol, and isobutanol) and 5 derived butyrate esters (ethyl butyrate, propyl butyrate, isopropyl butyrate, butyl butyrate, and isobutyl butyrate) from glucose (Figure 4A).

**Figure 4:**
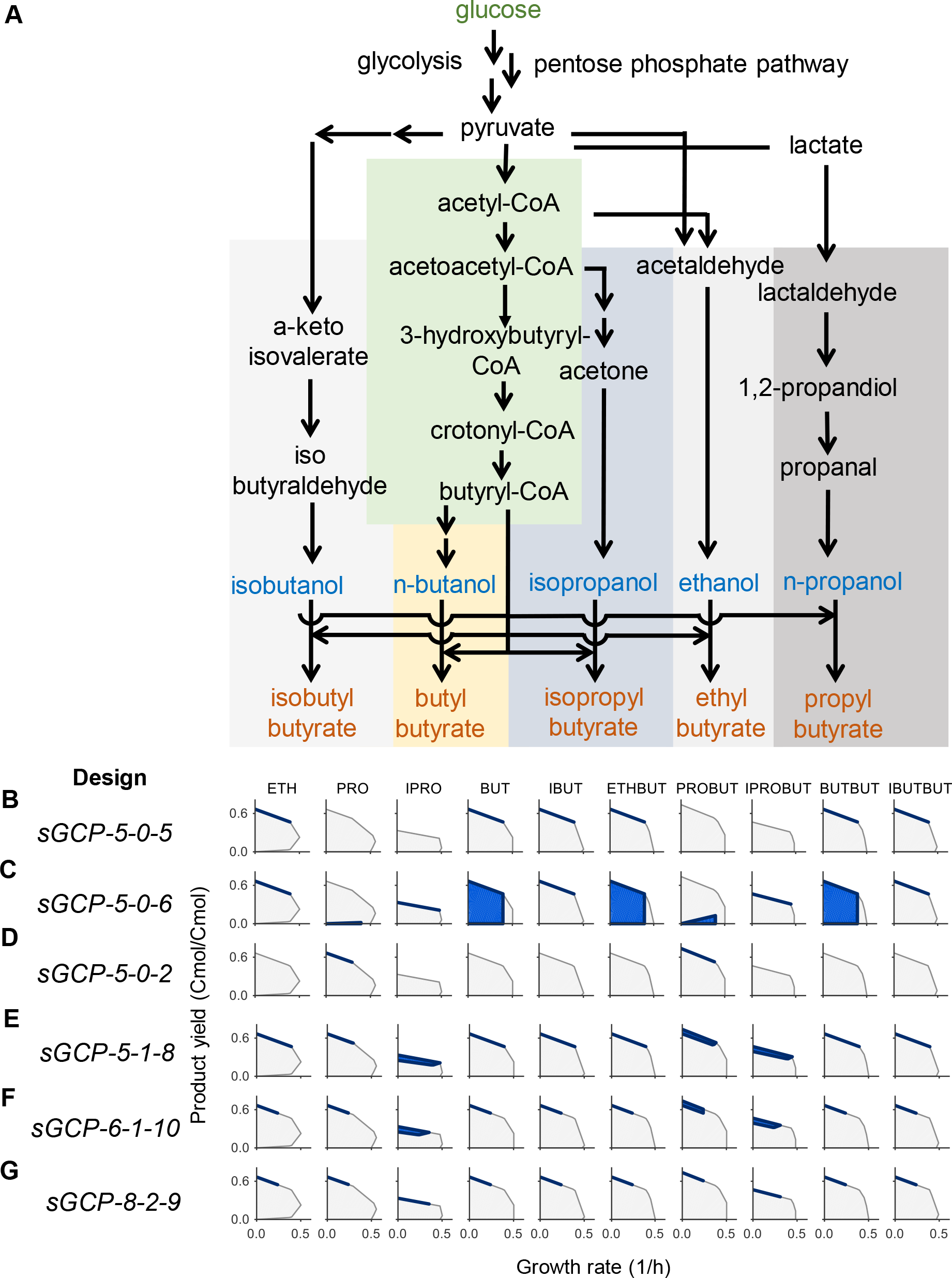
The 2-D metabolic phenotypic spaces of different *sGCP* designs using the core metabolic model. **(A)** Metabolic map, **(B)** *sGCP-5-0-5* design, **(C)** *sGCP-5-0-6* design, (D) *sGCP-5-0-2* design, **(E)** *sGCP-5-1-8* design, **(F)** *sGCP-6-1-10* design, and **(G)** *sGCP-8-2-9* design. Foreach panel, the gray and blue areas correspond to the phenotypic spaces of the wildtype and the optimal production strain, respectively.

In the first case study, we fixed the reaction module, i.e. *β*_*k*_ = 2 for ethanol dehydrogenase (FEM5) and ethanol export reaction (TRA1), in ModCell2 to emulate the same input as MODCELL (Supplementary File S3). The results showed that ModCell2 generated all the designs with the same *sGCP* objective values like MODCELL (Figure 4B, 4C, 4D) together with other alternative solutions (Supplementary File S3). Interestingly, ModCell2 only required 5 and 6 reaction deletions as opposed to 7 and 7 for the *sGCP-5-0-5* and *sGCP-5-0-6* designs, respectively. By setting the maximum reaction deletions to *α* ≥ 6, ModCell2 could find better design solutions with fewer deletion reaction requirement and higher objective values (Supplementary File S3).

In the second case study, we used the same model configuration but did not enforce the module reactions. By setting *α* = 5 and *β*_*K*_ = 1, we found the *sGCP-5-1-8* design that is compatible with all products and achieves the same objective values for products found in *sGCP-5-0-5*, *sGCP-5-0-6*, and *sGCP-5-0-2* (Figure 4E). The desirable phenotypic spaces can be further constrained for many products if *α* is increased from 5 to 6 (Figure 4F). Remarkably, by setting *α* = 8 and *β*_*k*_ = 2, we found a utopia point design, *sGCP-8-2-9*, without any trade-off among design objectives (Figure 4G). This utopia point design could not be achieved with *α* < 8 regardless of any *β*_*k*_ value.

Overall, the results demonstrate that ModCell2 can efficiently compute the Pareto front of modular cell designs. It can find better designs with fewer reaction deletion and module reaction requirements, improve design objective values, and enhance compatibility.

#### ModCell2 can identify designs with more compatibility than the conventional singleproduct designs

To evaluate if the conventional, single-product design strategy is suitable for modular cell engineering, we first used OptKnock to generate *wGCP* designs for the same 10 target molecules independently with various allowable reaction deletions (*α* = 2, 3, …,7). Likewise, we employed ModCell2 to produce *wGCP* designs using the same *α* and various β. To directly compare OptKnock and ModCell2 solutions, we calculated the *wGCP* design objective values for all products based on each OptKnock solution (Supplementary File S3). As expected, our result showed that ModCell2 and OptKnock designs have the same highest objective values for each product (Figure 5A). However, several OptKnock solutions were always dominated by ModCell2 solutions in all parameter configurations (Figure 5B). With *α* ≥ 4, ModCell2 could identify *wGCP-α-1* designs with the maximum compatibility of 10, while the best OptKnock designs only achieved the highest compatibility of 5 (Figure 5C, 5D).

**Figure 5:**
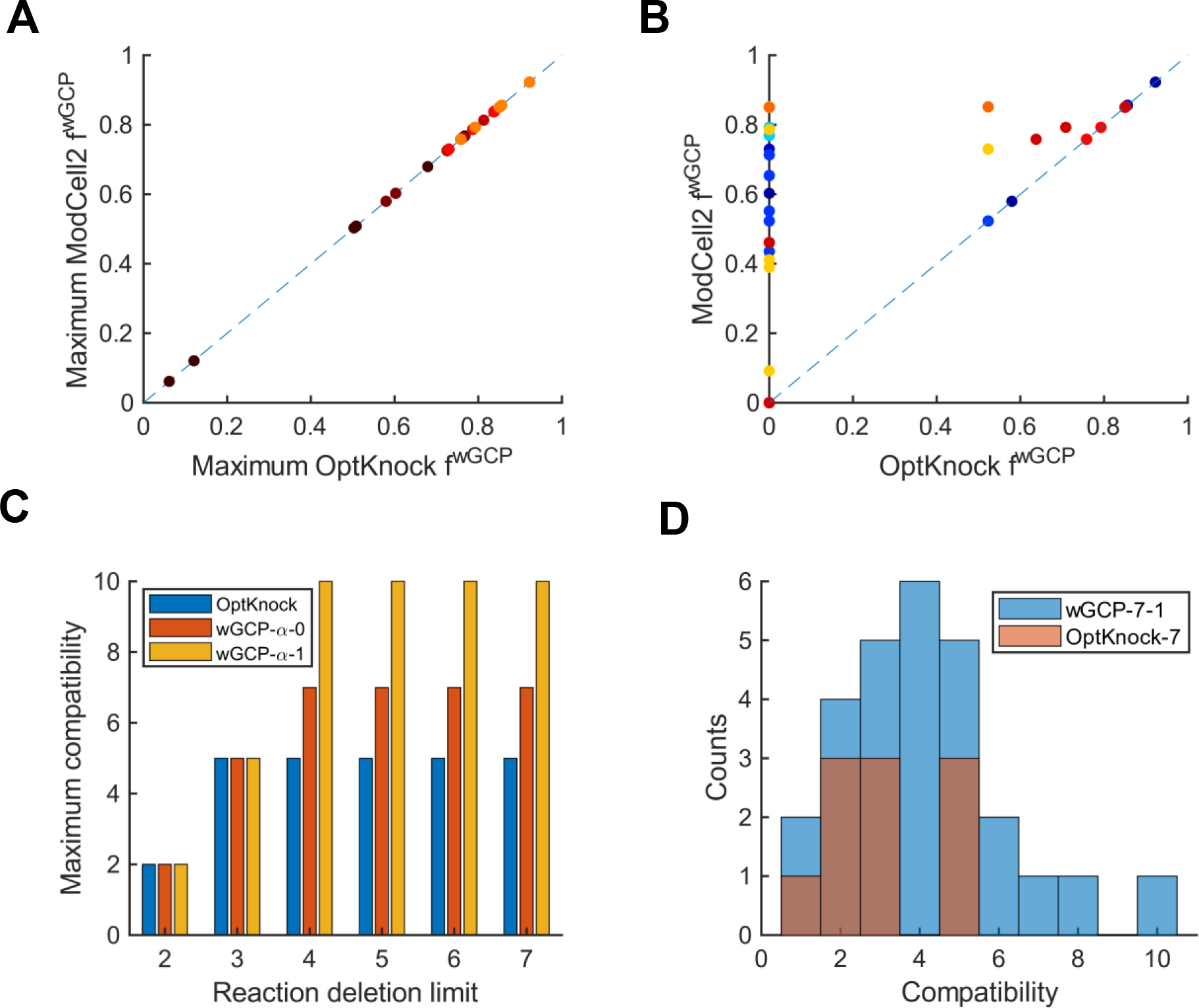
Comparison of strain design by OptKnock and Modcell2. (**A**) A correlation between the maximum objective values for each product generated by OptKnock and the equivalent values attained by ModCell2. Each point is colored based on the number of reaction deletions, with warmer colors corresponding to more reaction deletions. (**B**) A comparison between the Optknock objective vectors with at most 7 reaction deletions and the representative ModCell2 objective vector, *wGCP-7-1* which dominates them. Each color circle represents a pair of dominating *wGCP* design and dominated OptKnock solution (Supplementary File S3). (**C**) Maximum compatibility of OptKnock designs (blue), *wGCP* designs (β_k_ = 0, orange), wGCP designs (β_k_ = 1, yellow). (**D**) Compatibility distribution of Optknock (α = 7, orange) and *wGCP-7-1* (blue).

Overall, ModCell2 can generate modular cells compatible with the maximum number of modules and achieve high objective values. Single-product designs might not be compatible with a large number of products, and the solutions might be far from Pareto optimality.

### Exploring emergent features of modular cell design using an *E. coli* genome-scale network

#### Modcell2 can design modular cells using a large-scale metabolic network

To demonstrate that ModCell2 can be applied for a genome scale metabolic network, we tested it to generate *wGCP* designs for 20 target molecules with *α* = 4 and various (Supplementary File S4). The *α* value was chosen because with 4 deletions, OptKnock could identify single product designs with objectives above 60% of the theoretical maximum (Supplementary File S5). With *β*_*k*_ = 0, ModCell2 could identify modular cell designs with compatibility of 17, for example, the *wGCP-4-0-50* design featuring deletion of ACALD (acetaldehyde dehydrogenase, *adhE*), ACKr/PTAr (acetate kinase, *ack*; phosphotransacetylase, *pta*), GLYAT (glycine C-acetyltransferase, *kbl*), and LDH_D (lactate dehydrogenase, *ldhA*) (Figure 6A, 6D, Supplementary File S4). By analyzing all *wGCP-4-β_k_-d* designs (257 total for *β*_*k*_ = 0, 1, 2, and 3), we found that the ethanol and D-lactate production modules are most compatible with all modular cell designs (Figure 6A, 6C, Supplementary File S4). Among reaction deletions, *LdhA* (86% of designs), *Pta* (38%), and *AdhE* (25%) are the most frequent deletion reactions (Figure 6B). This finding is consistent with a comprehensive survey of metabolic engineering publications (55) showing that these deleted reactions appeared in most of *E. coli* engineered strains for production of fuels and chemicals. The result supports the potential use of modular cell engineering to systematically build modular platform strains.

**Figure 6:**
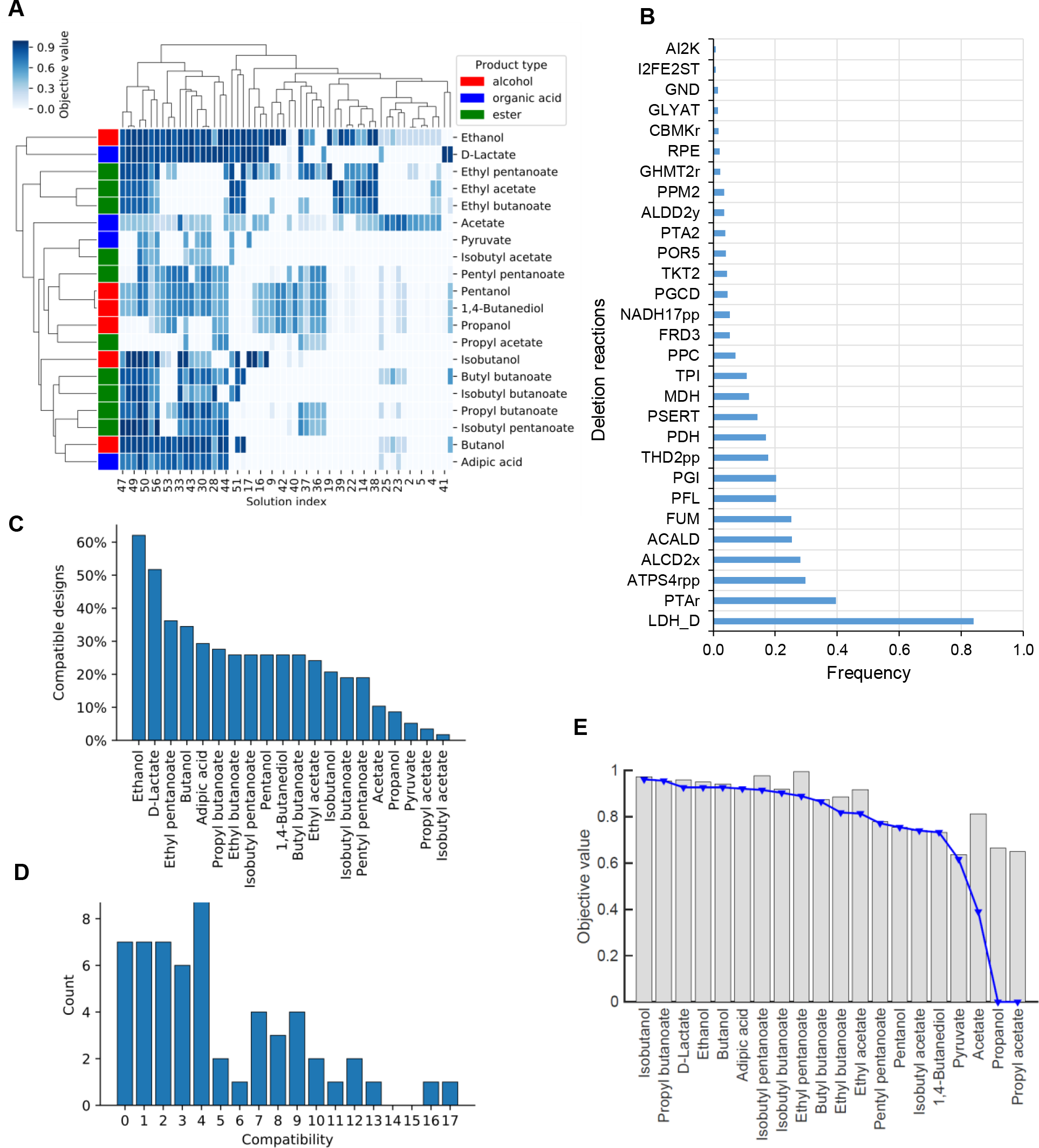
**(A)** Pareto front of *wGCP-4-0-d.* The columns correspond to different designs labeled by their design index, *d*, where the rows correspond to different products. **(B)** Frequency of the top deletion reactions. **(C)** Product compatibility distribution across designs. **(D)** Design compatibility. **(E)** Tradeoff between modularity and performance. The bars correspond to the maximum objective values attainable for each product whereas the blue line represent the objective values of the *wGCP-4-0-48alternative* design.

#### ModCell2 designs can capture combinatorial characteristics of production modules

To evaluate whether ModCell2 could capture the combinatorial properties among production modules, we analyzed the Pareto front of *wGCP-4-0-d* that have a total of 58 designs. Hierarchical clustering of this Pareto front revealed certain products with similar objective values across solutions, such as ethyl esters and butyrate esters (Figure 6A). These products together were compatible with different modular cells and exhibited metabolic similarity in their production modules. Thus, ModCell2 could generate designs that capture the combinatorial properties useful for modular cell engineering.

#### ModCell2 can identify highly compatible modular cells

Analysis of compatibility shows that certain modular cells can couple with production modules that may not exhibit the combinatorial properties (Figure 6D). However, there exists a tradeoff between the number of feasible designs and degree of compatibility. Some modular cell designs are compatible with up to 17 out of 20 products, for instance, the most compatible design, *wGCP-4-0-48*, featuring deletions of ACALD (*adhE*), ACKr/PTAr (*ack, pta*), GND (phosphogluconate dehydrogenase, *gnd*) and LDH_D (*ldhA*) (Supplementary File S4). An alternative design *wGCP-4-0-48-alternative* also exists where deletion of G6PDH2r (gluose-6-phosphate dehydrogenase, *zwf*) is replaced by that of GND, the first step in the oxidative pentose phosphate pathway. The gene deletions in the design *wGCP-4-0-48-alternative* are a subset of the modular *E. coli* strain TCS095, whose modular properties have recently been validated experimentally (14).

To determine if modular cell design is a viable alternative to single-product design, we also analyzed a potential tradeoff between design performance and modularity by comparing the maximum value of each objective across all solutions in the Pareto front and the single-product design optima. If production modules exhibit competing phenotypes, a modular cell will not achieve the same performance in all modules as a single-product design strain. Analysis of the most compatible design *wGCP-4-0-48-alternative* showed that it could achieve objectives within 4% of the single-product optima in 14 products and within 10% in 3 products (Figure 6E). This result indicates that it is feasible to identify highly compatible modular cell designs without a significant tradeoff between performance and modularity.

#### Analysis of potential tradeoff between robustness and modularity can identify conserved metabolic features

To evaluate the robustness of modular cells, we analyzed the compatibility change (*CD*) of *wGCP-4-0* designs with compatibilities of 4 or greater (Figure S3 in Supplementary File 2). Remarkably, the result shows that only 7.5% of potential reaction deletions were detrimental to the robustness of modular cells while the large remaining portion did not affect *CD* values. Out of the 85 reactions whose deletion affected compatibility, only a few appeared consistently across the designs. For instance, deletion of TPI (triose-phosphate isomerase, *tpi*) led to an average compatibility loss of 95%, inactivating most modular cell designs. Based on flux variability analysis, TPI must operate in the forward direction by converting glycerone phosphate (dhap) to glyceraldehyde-3-phosphate (g3p) to drive sufficient flux through glycolysis and hence preventing synthesis of undesired byproducts (D-lactate or 1,2-propanediol) from dhap. Likewise, deletion of carbon dioxide and water transport and exchange reactions caused compatibility loss across all designs. Pyruvate carboxylase (PPC) is an important reaction to channel carbon flux through the Krebs cycle (56), and hence, deletion of PPC reduces compatibility in most modular cell designs with an average *CL* of 43%.

While some reaction deletions are critical for modular cell robustness, others are associated with specific products. For example, deletion of PDH (pyruvate dehydrogenase complex, *lpd/aceEF*) removes compatibility in all butanol-derived designs, indicating PFL (pyruvate formate lyase, *pfl*) is not an appropriate route. To make heterologous butanol-derived molecules under anaerobic conditions, FDH (NADH-dependent formate dehydrogenase, *fdh*) is required in butanol-derived modules where enzymatic reaction pairs of PFL and FDH could substitute PDH known to be anaerobically inhibited.

Overall, analysis of tradeoff between modularity and robustness can identify not only the conserved metabolic features of modular cells but also potential bottlenecks in specific production modules.

#### Enabling metabolic switch among different design objectives using ModCell2

The ability to dynamically control growth and production phases can potentially enhance product titers, rates, and yields. For instance, two-phase fermentation can be employed where growth phase is optimized for biomass synthesis and stationary phase for chemical production (57). Using ModCell2, we investigated the feasibility to design optimal strains to toggle switch desirable production phenotypes.

To design a *wGCP→NPG* metabolic switch, we first used our reference *wGCP* design as a parent strain (Figure 7A) and then employed ModCell2 to identify the most compatible *wGCP→NPG* designs. With 5 additional deletions, we could find *wGCP→NPG* designs that encompass both *wGCP* and *NPG* phenotypes, for instance, the *sup-NGP-5-0-23* design featuring deletion of PGI (glucose-6-phosphate isomerase, *pgi*), MDH (malate dehydrogenase, *mdh*), ASPT (L-aspartase, *aspT*), Tkt2 (transketolase, *tktB*), and ATPS4rpp (ATP synthase, *atp*) (Figure 7B). The deletion reactions in the *wGCP→NPG* designs appear in both catabolic (PGI, ATPS4tpp) and anabolic (ASP, TKT2) processes, responsible for growth disruption and direction of carbon flow to the biosynthesis of target products.

**Figure 7:**
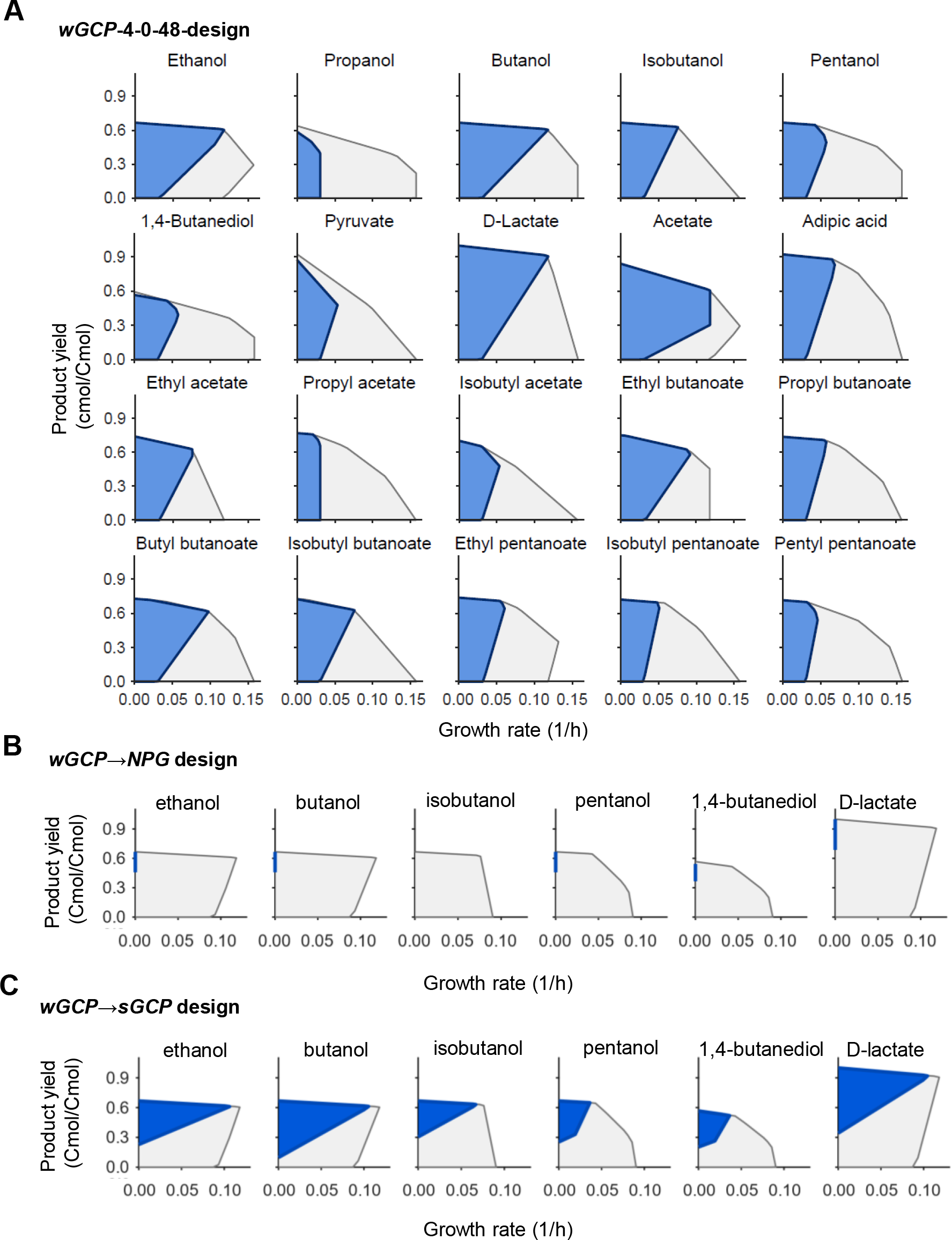
Production phenotypes of (**A**) the wild type (gray) and the representative, highly-compatible design *wGCP-4-0-48-alternative* (blue), (**B**) the *wGCP→NPG* design, *sup-NGP-5-0-23*, and (**C**) the *wGCP→sGCP* design, *sup-sGCP-6-0-39*.

Likewise, we used ModCell2 to design a *wGCP→sGCP* metabolic switch. We identified the most compatible *wGCP→sGCP* designs with 6 additional deletions, for instance, the *sup-sGCP-6-0-39* design featuring the deletion of MGSA (Methylglyoxal synthase, *mgsA*), ALCD2x (alcohol dehydrogenase, *adhE*), PFL, MDH, FADRx (FAD reductase, *fadI*), and GLUDy (NADP+dependent glutamate dehydrogenase, *gdhA*) (Figure 7C). Different from the *wGCP→NGP* metabolic switch, all deletions in the *wGCP→sGCP* designs are involved the elimination of biosynthesis pathways of undesirable byproducts.

While it is feasible to metabolically switch among different production phenotypes, it not only requires more reaction deletions but also reduces the product compatibility. For instance, the *wGCP→sGCP* and *wGCP→NGP* designs are only compatible with 5 products while the *wGCP* parent design have a compatibility of 17 out of 20 products with 4 deletions. The main reason is that both *wGCP→sGCP* and *wGCP→NGP* designs must eliminate all possible redundant pathways that result in biosynthesis of undesirable byproducts.

## CONCLUSION

In this study, we developed a multiobjective strain design platform for modular cell engineering. With a new developed algorithm and computational platform, ModCell2 enables flexible design of modular cells that can couple with production modules to exhibit desirable production phenotypes. In comparison to the first-generation strain design platform, ModCell2 can handle large-scale metabolic networks and identify better solutions that require fewer genetic modifications and exhibit more product compatibility. Different from the conventional single product strain design, ModCell2 can find solutions that are Pareto optimal with negligible tradeoffs among modularity, performance, and robustness. We envision ModCell2 is a useful tool to implement modular cell engineering and fundamentally study modular designs in natural and synthetic biological systems.

## ACKNOWLEDGEMENTS

This research was financially supported in part by the NSF CAREER Award
(NSF#1553250) and the DOE subcontract Grant (DE-AC05-000R22725) by the Center of Bioenergy Innovation (CBI), the U.S. Department of Energy Bioenergy Research Center funded by the Office of Biological and Environmental Research in the DOE Office of Science. The funders had no role in the study design, data collection and analysis, decision to publish, or preparation of the manuscript.

